# Dietary Induction and Modulation of Ferroptosis in *Caenorhabditis elegans*

**DOI:** 10.1101/772079

**Authors:** Marcos A. Perez, Leslie Magtanong, Scott J. Dixon, Jennifer L. Watts

## Abstract

Ferroptosis is an iron-dependent form of regulated cell death associated with oxidized polyunsaturated phospholipids. Understanding the role of this process in vivo has been slowed by the lack of readily accessible model systems. Exposing the nematode *Caenorhabditis elegans* to the polyunsaturated fatty acid dihomogamma-linolenic acid (DGLA; 20:3n-6) causes germ cell death and sterility that is largely independent of the canonical apoptosis pathway. Here we demonstrate that DGLA-induced germ cell death is modulated by small molecule ferroptosis inhibitors, genetic manipulation of ferritin, NADPH oxidase, and glutathione peroxidases, and by dietary co-supplementation with oleic acid. Thus, DGLA-induced germ cell death in *C. elegans* is highly analogous to ferroptosis in mammalian cells. DGLA can also induce ferroptosis in human cells, further highlighting this omega-6 PUFA as a metabolic instigator of ferroptosis. Together, these results establish *C. elegans* as a powerful animal model to study the induction and modulation of ferroptosis by dietary fats.

**Highlights:**

- Dietary dihomogamma-linolenic acid (DGLA)-induced germ cell death in *C. elegans* is alleviated by small molecule antioxidants and iron chelators
- Dietary and endogenous oleic acid protects from DGLA-induced ferroptosis
- Ether-lipid deficiency increases sensitivity to DGLA-induced ferroptosis
- DGLA specifically induces ferroptosis in human cancer cells

## INTRODUCTION

Long-chain polyunsaturated fatty acids (PUFAs) are essential in the diet of humans, although there is still debate regarding the optimal amounts of various species of dietary fats (Mukhopadhyay, 2012). PUFAs are classified as omega-6 (n-6) or omega-3 (n-3), depending on the position of the terminal double bond in the molecule. The omega-6 fatty acids are precursors for powerful signaling molecules, such as prostaglandins and leukotrienes that promote inflammatory responses. These responses are important for fighting pathogens and for normal reproduction, but excess omega-6 fatty acids are associated with disease states, such as cardiovascular disease and cancer (Harris et al., 2009).

Ferroptosis, an iron-dependent form of regulated non-apoptotic cell death, is defined by three conserved hallmarks – the presence of oxidized polyunsaturated fatty acids (PUFAs), the availability of redox-active iron, and defective or inhibited lipid peroxide repair (Dixon and Stockwell, 2019). Ferroptosis has been demonstrated in several mammalian systems, including various cancer cell lines (Dixon et al., 2012; Dixon et al., 2014), inducible *Gpx4*^−/−^ mutant mice (Friedmann Angeli et al., 2014; Ingold et al., 2018), rat hippocampal slices and oligodendrocyte cultures (Dixon et al., 2012; Skouta et al., 2014). In mammalian cells, ferroptosis can be suppressed by small molecule iron chelators and radical trapping antioxidants (Stockwell et al., 2017). The physiological relevance and regulation of ferroptosis is currently a topic of great interest, but the influence of diet on ferroptosis has not been examined. Progress on this and other related questions has been hindered by the absence of accessible and easy-to-manipulate animal models of the ferroptotic process.

The nematode *Caenorhabditis elegans* (*C. elegans*) is a powerful model for investigating the importance of dietary fatty acids in the development and maintenance of the germ line. This species synthesizes a wide range of fatty acids de novo (Hutzell and Krusberg, 1982; Watts and Browse, 2002), and grows and reproduces over a wide range of temperatures. Reproduction is an energetically intensive process, where at peak egg laying an adult hermaphrodite converts the equivalent of its entire body mass to eggs daily (Hirsh et al., 1976). Dietary fats are an important source of lipids for *C. elegans* egg production, and fatty acids in the body turn over every 24 hours (Dancy et al., 2015).

Previously, we discovered that *C. elegans* cultured in the presence of the polyunsaturated fatty acid dihomogamma-linolenic acid (DGLA, 20:3n-6) became sterile (Watts and Browse, 2006). Wild-type hermaphrodites raised on DGLA are severely deficient in germ cell production, and exposure to lower levels of DGLA which did not cause 100% sterility in the population nevertheless led to reduced numbers of germ cells, sperm, and progeny (Watts and Browse, 2006). Genetic pathways regulating oxidative stress responses, lipid homeostasis, and lifespan modulated the sensitivity to dietary DGLA (Watts and Browse, 2006; Webster et al., 2013), as did the manipulation of PUFA oxidation enzymes (Deline et al., 2015).

In this study we tested the hypothesis that dietary DGLA-induced germ cell death occurs through ferroptosis. Using chemical biology, genetic and lipidomic approaches, we report that iron, reactive oxygen species (ROS) and lipid metabolism modulate the effects of dietary DGLA on the worm reproductive system. Furthermore, DGLA supplementation is sufficient to induce ferroptosis in human cancer cells, demonstrating the possibility of diet-mediated ferroptosis across species.

## RESULTS

### Dietary DGLA triggers ferroptotic germ cell death

Dietary DGLA is a potent inducer of germ cell death in *C. elegans*, with genetic evidence suggesting this could be in part an oxidative process (Deline et al., 2015; Watts and Browse, 2006; Webster et al., 2013). To test whether this process involved ferroptosis, we co-treated wild-type animals with dietary DGLA and the ferroptosis-specific lipophilic radical trapping antioxidant ferrostatin-1 (Fer-1) (Dixon et al., 2012; Zilka et al., 2017). Strikingly, in animals co-treated with DGLA and Fer-1, both germ cell death and sterility were reduced compared to worms treated with DGLA only (Figure 1A). Fer-1 alone had no effect of fertility. Not only were there more fertile worms in the DGLA + Fer-1 treatment group compared to DGLA alone, but the number of fertile worms with normal gonad development, as assayed by DAPI staining, was increased compared to the control group (Figures 1B,C).

**Figure 1.**
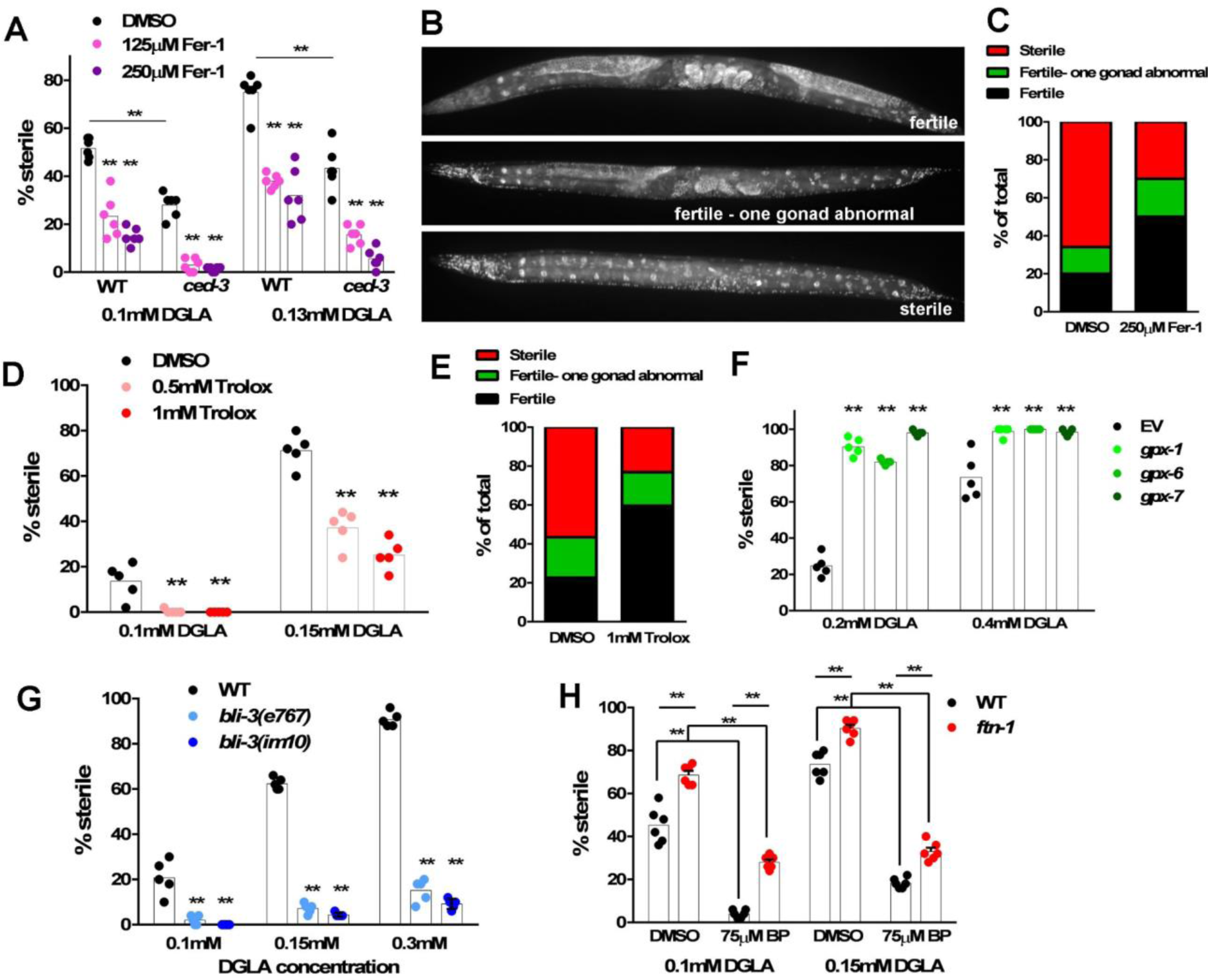
Antioxidants, glutathione peroxidase, NADPH oxidase and cellular iron modulate DGLA-induced germ cell death. (A) Percentage (%) sterility of wild-type and *ced-3 C. elegans* raised on 0.1 mM or 0.13 mM DGLA and two doses of ferrostatin-1 (Fer-1). (B) Representative images of wild-type *C. elegans* after exposure to 0.1 mM DGLA. Fertile worms may have abnormal (diminished) anterior or posterior gonads. (C) Quantification of fertility phenotypes based on the categories in (B) in a sample of 50 wild-type worms treated with 0.1mM DGLA ± 250 μM Fer-1. (D) % sterility of wild-type worms exposed to DGLA ± Trolox (vitamin E). (E) Quantification of fertility phenotypes based on the categories in (B) in a sample of 50 wild-type worms treated with 0.15mM DGLA ± 1mM Trolox. (F) % sterility of RNAi knockdown of the phospholipid glutathione peroxidase genes *gpx-1, gpx-6*, and *gpx-7*. (G) % sterility of wild-type or NADPH oxidase/duox (*bli-3*) mutant worms after dietary exposure to DGLA. (H) % sterility of wild-type or *ftn-1* mutant worms exposed to dietary DGLA ± 2,2’ bipyridine (BP). In (A, D, and F-H), each point represents an independent experiment comprising 50 worms for each treatment. Statistical significance was determined by two-way ANOVA with a Tukey’s test for multiple comparisons, two asterisks represent P-values <0.0001 when compared with control worms. All experiments used *E. coli* OP50 bacteria, except for (D), which used *E. coli* HT115 for expression of the RNAi feeding constructs. The addition of DMSO to plates for small molecule supplementation, as well as the *E. coli* food source (OP50 or HT115) affected the uptake of DGLA, necessitating the use of different DGLA concentrations for different experiments. DGLA uptake as measured by gas chromatography-mass spectrometry (GC-MS) in nematodes is reported in Supplemental Table 1.

We previously demonstrated that dietary DGLA led to germ cell death accompanied by an increase of apoptotic corpses, and that sterility was reduced, although not eliminated, in mutants in the core apoptotic pathway (Watts and Browse, 2006; Webster et al., 2013). This implied that while the apoptotic pathway contributes to DGLA-induced cell death, another regulated cell death pathway may contribute to the germ cell depletion observed in apoptosis-deficient *ced-3* and *ced-4* mutants. Indeed, Fer-1 suppressed sterility in *ced-3* mutants treated with DGLA, indicating that ferroptosis and apoptosis are both activated in response to dietary DGLA (Figure 1A). The lipophilic vitamin E derivative Trolox also suppressed the sterility in DGLA-treated worms, indicating that this protective effect was likely linked to inhibition of lipid ROS formation (Figure 1D). DAPI staining of DGLA + Trolox-treated worms displayed increased numbers of fertile worms with normal gonad development compared to DGLA only-treated worms (Figure 1E). Together, these results demonstrate that known small molecule ferroptosis inhibitors reduce germ cell death induced by dietary DGLA in *C. elegans*, in parallel to the classic apoptotic machinery.

### DGLA-induced sterility is modulated by redox and iron metabolism

The glutathione-dependent phospholipid hydroperoxidase GPX4 inhibits ferroptosis in mammalian cells (Yang et al., 2014). *C. elegans* contains eight glutathione peroxidase orthologs, *gpx-1* through *gpx-8* (Sakamoto et al., 2014). Four *gpx* genes show the greatest similarity to phospholipid glutathione peroxidases, in that they all contain selenocysteine in the active site (Ingold et al., 2018), but sequence similarity is not high enough to identify a single definitive GPX4 homolog. We therefore used RNAi knockdown to deplete the expression of three *gpx* genes, *gpx-1, gpx-6*, and *gpx-7*. We found that knockdown of each of these genes led to increased susceptibility to sterility induced by dietary DGLA (Figure 1F). Thus, as in mammals (Ingold et al., 2018), phospholipid glutathione peroxidases protect *C. elegans* germ cells from ferroptosis induced by dietary fatty acids.

In mammalian cells, NADPH oxidase (NOX) is an important source of cellular ROS that can promote ferroptosis in some contexts (Dixon et al., 2012; Xie et al., 2017). *bli-3* encodes a dual oxidase (Duox) enzyme that is the sole *C. elegans* NOX homolog. The *bli-3(e767)* strain carries a point mutation (G246D) in the peroxidase domain, while the *bli-3(im10)* strain carries a mutation in the NOX domain (P1311L) (Moribe et al., 2012). When exposed to dietary DGLA, both *bli-3(e767)* and *bli-3(im10)* worms exhibited strong suppression of sterility (Figure 1G).

In addition to the presence of oxidized PUFAs and modulation by endogenous and exogenous antioxidants, a hallmark of ferroptosis is the accumulation of redox-active iron (Stockwell et al., 2017). Intracellular iron is bound to ferritin, which controls iron release in a non-toxic manner. Consequently, *ftn-1* mutants have elevated levels of intracellular iron (Valentini et al., 2012). We observed that the ferritin mutant *ftn-1* is more susceptible to dietary DGLA-induced cell death than wild-type worms (Figure 1H). Furthermore, treating both wild-type and *ftn-1* worms with the iron chelator 2,2’ bipyridine, which reduces cellular iron levels, suppressed the sterility induced by dietary DGLA (Figure 1H). Together, these experiments demonstrate that DGLA-induced germ cell ferroptosis in *C. elegans* is modulated by redox and iron metabolism, much like ferroptosis in mammalian cells induced by glutathione depletion or GPX4 inhibition.

### Dietary and endogenous monounsaturated fatty acids inhibit germ cell ferroptosis

In mammalian cells, the addition of oleic acid (OA), a monounsaturated fatty acid (MUFA), protects HT-1080 cells from ferroptosis by lowering the levels of PUFAs in cellular membranes (Magtanong et al., 2019). To test whether exogenous OA could protect *C. elegans* germ cells from dietary DGLA-induced sterility in an analogous manner, we fed wild-type worms various combinations of OA and DGLA. Similar to mammalian cells, we observed strong reduction of dietary DGLA-induced sterility when OA was included in the culture medium (Figure 2A). Gas chromatography-mass spectrometry (GC-MS) analysis of worm fatty acid composition indicated that dietary OA was incorporated into worm lipids, increasing the relative levels of OA compared to vehicle-treated worms. Levels of 18:2n-6 (LA) and 20:5n-3 (EPA) also increased, but relative levels of other C20 PUFAs, including DGLA and AA, were lowered when OA was added to the growth media (Figure 2C and Supplementary Table 1). This indicates that dietary OA can protect cells from ferroptosis by replacing DGLA and AA in cellular membranes.

**Figure 2.**
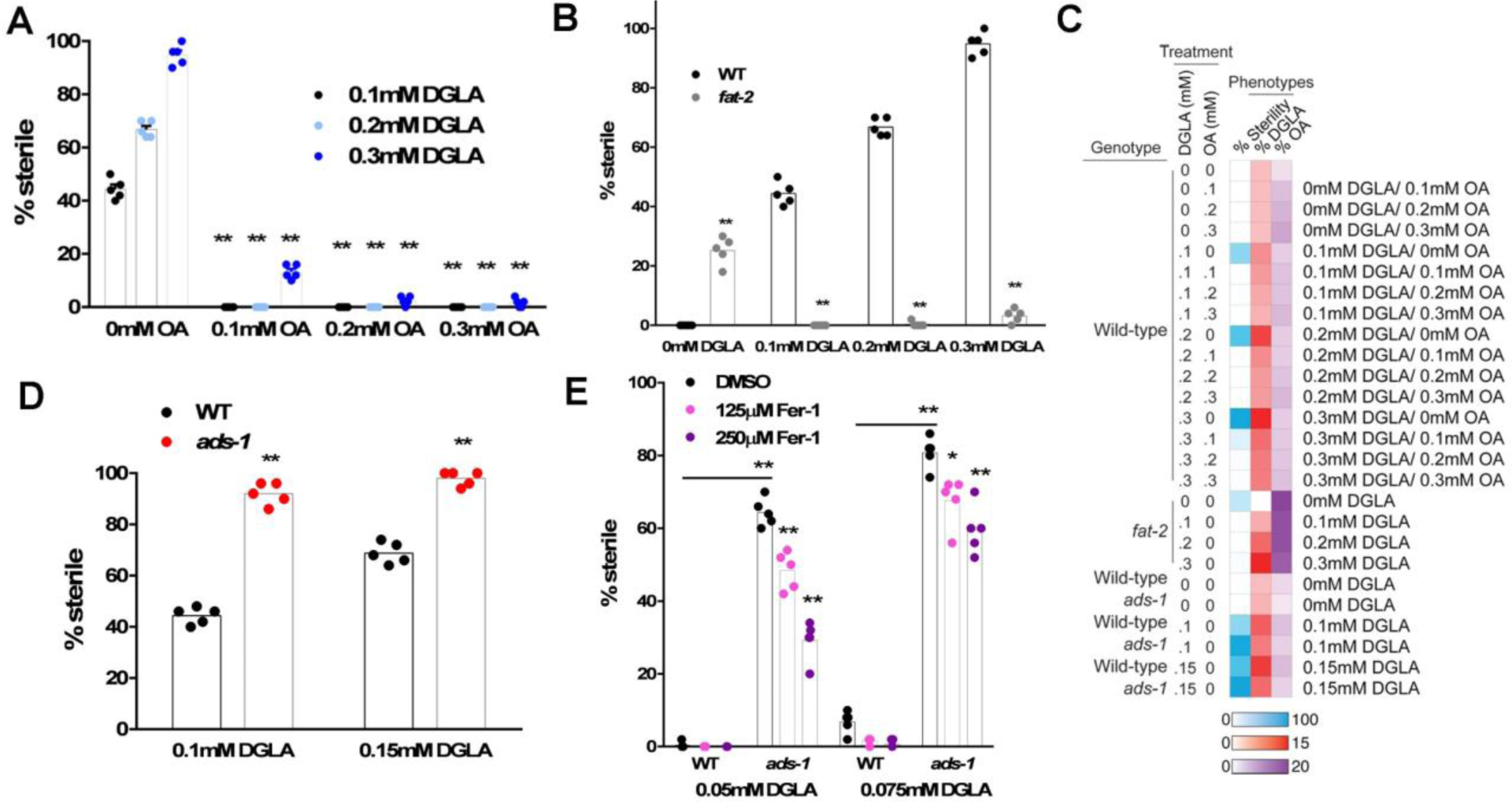
Dietary and endogenous oleic acid and ether lipids protect from dietary DGLA-induced germ cell death. (A) Percentage (%) sterility of wild-type worms fed the indicated combinations of oleic acid (OA) and DGLA. Graphs show the average percentage of sterility in five populations of 50 worms exposed to various concentrations of DGLA and OA. Asterisks show significance P<0.0001) of various OA treatments compared to the 0 OA controls. (B) % sterility of wild-type and *fat-2* mutants raised on DGLA plates. (C) Heat map showing the % sterility, % DGLA composition in worm extracts, and % OA composition in worm extracts for experiments shown in panels 2A, 2B, and 2C. (D) % sterility of wild-type and *ads-1* mutants raised on DGLA. (E) % sterility of wild-type and *ads-1* mutants raised on low doses of DGLA in the presence and absence of two doses of Fer-1. For panels A, B, D, and E, significance was determined by two-way ANOVA with Tuckey’s multiple compassion test, * P<0.01, **P<0.0001.

The *fat-2* gene encodes a delta-12 fatty acid desaturase, and *fat-2* mutant worms are unable to convert OA to LA. Consequently, they accumulate high levels of OA in all lipids and lack C18 and C20 PUFAs (Watts and Browse, 2002). To test if endogenous OA protects from DGLA-induced sterility, we exposed *fat-2* mutants to dietary DGLA. Approximately 25% of *fat-2* worms are sterile in the absence of dietary DGLA due to lack of PUFAs required for efficient reproduction (Figure 2B). This sterility was rescued by dietary DGLA, and furthermore, the *fat-2* worms were almost completely protected from DGLA-induced germ cell death, even though the worms accumulated DGLA to levels that normally cause sterility in wild-type worms. This protection is associated with high levels OA in the corresponding worm lipids (Figure 2C and Supplementary Table 1). Thus, elevating endogenous MUFA levels is sufficient to prevent DGLA-induced germ cell ferroptosis. These results illustrate that both exogenous and endogenous MUFAs can inhibit germ cell ferroptosis in *C. elegans*.

### Ether lipid deficiency exacerbates DGLA-induced germ cell ferroptosis

The presence of ether-linked lipids, especially plasmalogens, in cells has been associated with protection against oxidative stress (Engelmann, 2004; Wallner and Schmitz, 2011). We previously found that mutation of genes required for ether lipid biosynthesis increases sensitivity to oxidative stress induced by tert-butyl peroxide or paraquat (Shi et al., 2016). The *C. elegans ads-1* gene encodes alkylglycerone phosphate synthase (AGPS), which catalyzes the rate-limiting step for ether lipid biosynthesis. We treated ether lipid-deficient *ads-1* mutant worms with dietary DGLA and found much higher levels of sterility than wild-type controls (Figure 2D), even though the uptake of DGLA was similar or slightly lower than in the wild-type controls (Figure 2C and Supplementary Table 1). Fer-1 treatment had no effect on the sterility of *ads-1* mutant worms grown on plates with 0.1 mM DGLA or 0.15 mM DGLA (data not shown). However, *ads-1* mutants grown on lower doses of DGLA, which caused very little sterility in wild-type animals, caused considerable sterility, and this sterility was reduced by the addition of Fer-1 (Figure 2E). Therefore, ether lipid synthesis protects *C. elegans* from dietary DGLA-induced ferroptosis.

### DGLA induces ferroptosis in human cancer cells

In mammalian cells, ferroptosis is typically induced and studied using synthetic small molecule inhibitors of glutathione metabolism or GPX4 function (Stockwell et al., 2017). Ferroptosis has been associated with the oxidation of two omega-6 PUFAs, arachidonic acid (AA, C20:4n-6) and adrenic acid (AdA, C22:4n-6), esterified to phosphatidylethanolamine (PE) (Kagan et al., 2017). Whether exogenous omega-6 PUFAs such as DGLA or AA are sufficient to induce ferroptosis as single agents in cancer cells is unclear. To test this hypothesis, we used a classic ferroptosis-sensitive cell line, HT-1080 fibrosarcoma, stably expressing nuclear-localized mKate2 (denoted HT-1080^N^). This enabled us to monitor cell death using the scalable time-lapse analysis of cell death kinetics (STACK) approach (Forcina et al., 2017). Much like a positive control small molecule inducer of ferroptosis, erastin2, exogenous DGLA (500 μM) was capable of triggering ferrostatin-1-sensitive cell death within 24 h (Figure 3A,B). At the same concentration, exogenous AA induced cell death, but this was not effectively suppressed over time by co-treatment with Fer-1, indicating that at this dose exogenous AA may be contributing to other modes of cell death (Figure 3A). At lower doses (250 μM and below), both DGLA and AA exhibited little lethality in HT-1080^N^ cells (Figure 3A). Thus, exogenous omega-6 PUFAs are sufficient to trigger ferroptosis as single agents in human cancer cells in a structure-specific manner.

**Figure 3.**
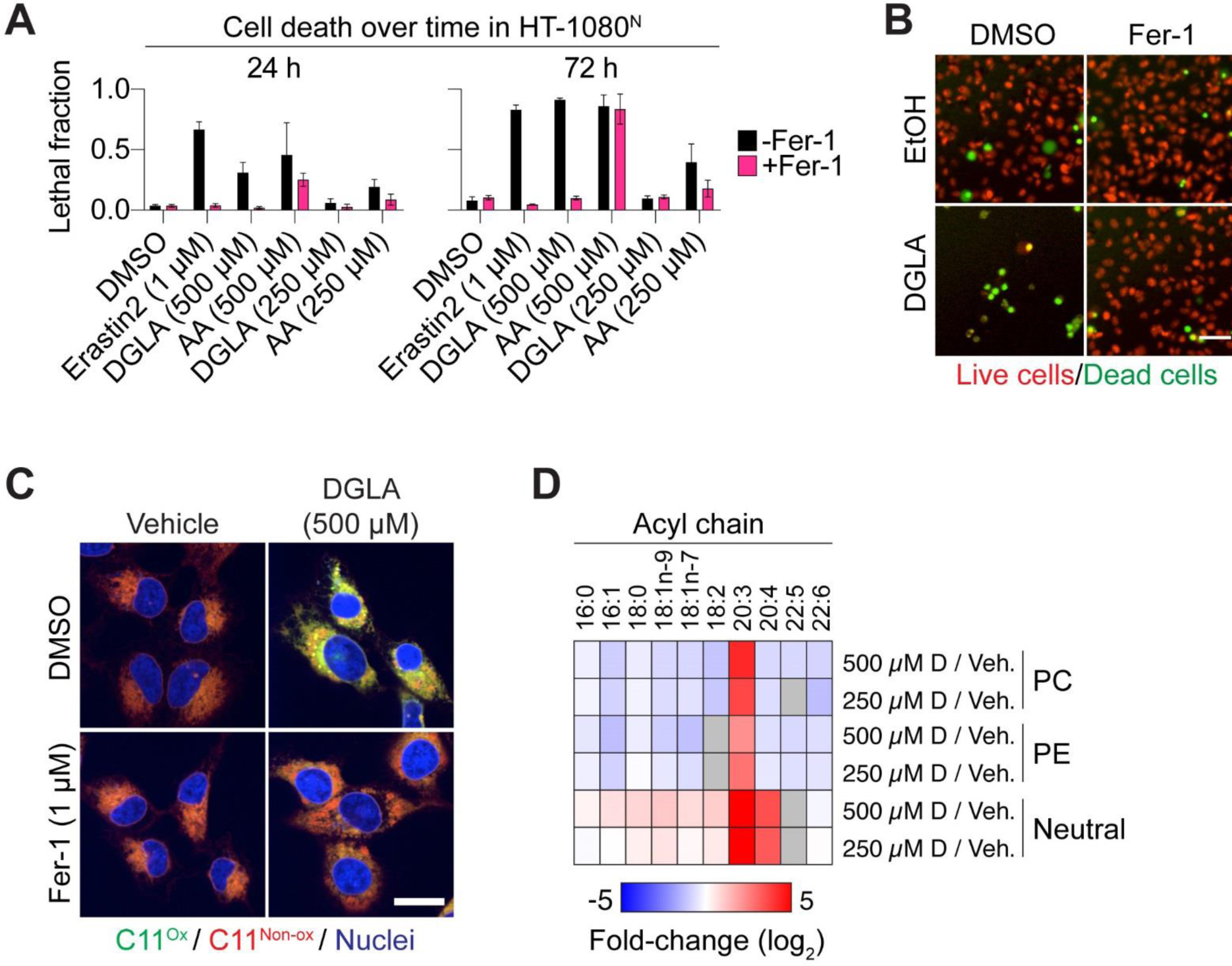
DGLA induces ferroptosis in human cells. (A) Cell death (lethal fraction) over time in HT-1080^N^ cells treated ± erastin2, DGLA or arachidonic acid (AA) and ± Fer-1 (1 μM). (B) Representative images showing a field of HT-1080^N^ cells treated ± DGLA (500 μM) ± ferrostatin-1 (Fer-1, 1 μM). Live cells express nuclear-localized mKate2; dead cells take up SYTOX Green. Images were acquired at 72 h post-treatment. (C) Confocal images of C11 BODIPY 581/591 (herein “C11”) in HT-1080 cells. After treatment for 8 h, cells were labeled with C11 (5 μM) and Hoechst (1 μg/mL) prior to imaging. C11^Ox^: Oxidized C11; C11^Non-Ox^: Non-Oxidized C11. Scale bar = 20 μm. (D) Heat map showing the relative fold changes in fatty acid composition of various lipid classes after 6 hours of growth with DGLA (500 μM or 250 μM) versus ethanol controls. PC: phosphatidylcholine, PE: phosphatidylethanolamine, neutral: neutral lipids. Grey boxes indicate situations with fatty acid concentrations at less than 0.4%. Data represent mean values from three independent experiments.

One hallmark of ferroptosis is the accumulation of membrane lipid ROS (Magtanong et al., 2019; Tarangelo et al., 2018). In mammalian cells lipid ROS can be effectively visualized using C11 BODIPY 581/591 (herein referred to as ‘C11’) together with confocal imaging (Magtanong et al., 2019). C11 is a ratiometric lipid peroxidation sensor that fluoresces red when it is not oxidized and green when it is oxidized (Pap et al., 1999). DGLA-treated (500 μM) HT-1080 cells exhibited a significant increase in membrane C11 oxidation, and this oxidation was completely blocked by co-treatment with Fer-1 (1 μM) (Figure 3C). These effects on lipid ROS accumulation were paralleled by changes in membrane phospholipid composition, as determined using thin-layer chromatography (TLC) and GC-MS. HT-1080^N^ cells treated with a lethal concentration of DGLA (500 μM, 6 h) exhibited a substantial increase in C20:3-containing neutral lipids (including free fatty acids, triacylglycerols, and diacylglycerols) and phospholipids, both phosphatidylcholines (PCs) and to a lesser extent phosphatidylethanolamines (PEs) (Figure 3D, Supplementary Table 1). DGLA-treated cells exhibited increased levels of AA in neutral lipids, consistent with conversion of C20:3 into C20:4. However, in phospholipids (both PC and PE) the levels of C20:4 and C22:5 and C22:6 were reduced, suggesting that DGLA ultimately displaces these species. Notably, higher levels of C20:3 incorporation into PCs, but not PEs, were observed at the lethal (500 μM) versus non-lethal (250 μM) DGLA concentration (Figure 3D). Furthermore, we observed relative decreases in all protective MUFA-containing PCs and PEs (Figure 3D). The combination of increased C20:3-PUFA-PLs and decreased MUFA-PLs may together yield membranes with a heightened sensitivity to excess lipid oxidation and the execution of ferroptosis.

## DISCUSSION

Here we report that ferroptosis can be induced in *C. elegans* germ cells exposed to the polyunsaturated fatty acid DGLA. This lethal phenotype is promoted by NOX activity and iron, and opposed by worm GPX4 orthologs and antioxidants. In these respects, the process observed in *C. elegans* germ cells is highly analogous to ferroptosis in mammalian cells (Table 1 and references therein). The specificity of DGLA-induced ferroptosis for the germ cells of young animals is of great interest. Dietary lipids may be preferentially concentrate in these cells during the rapid and metabolically demanding process of cell formation that occurs in the syncytial gonad. Another possibility is that the germline itself may be deficient in antioxidant defenses (or enriched in prooxidant enzymes) that render these cells hypersensitive to the accumulation of lipid peroxides compared to somatic cells. Interestingly, ferroptosis may also occur in intestinal cells of aged *C. elegans* (Jenkins, BioRxiv 2019). Here, the aging process itself may lead to the loss of control over iron homeostasis or other processes that normally restrain ferroptosis in somatic tissues. Together, these results establish *C. elegans* as a powerful and genetically tractable animal model of ferroptosis.

**Table 1.**
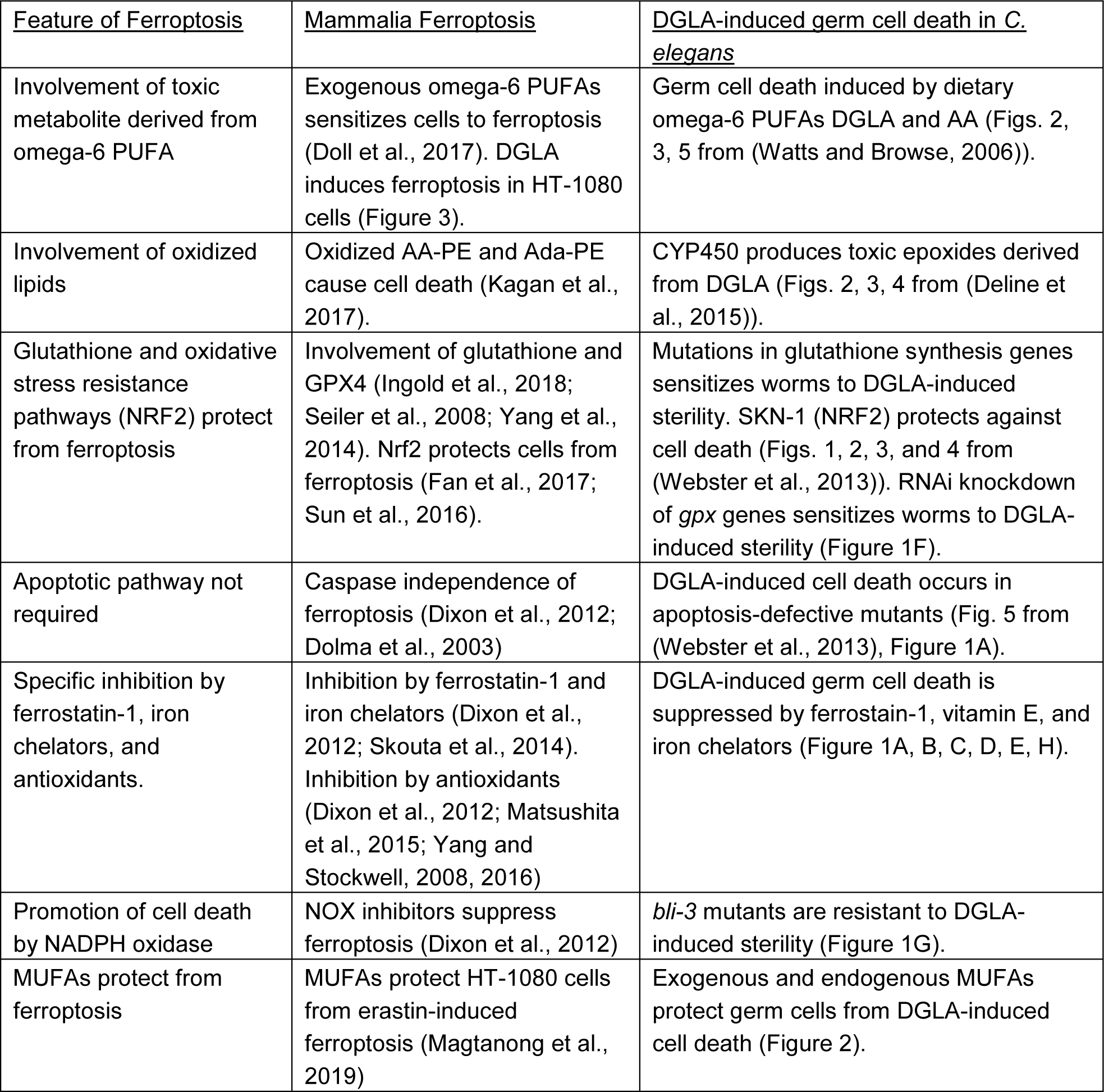
Common features of DGLA-induced germ cell death and mammalian ferroptosis.

| <u>Feature of Ferroptosis</u> | <u>Mammalia Ferroptosis</u> | <u>DGLA-induced germ cell death in *C. elegans*</u> |
| --- | --- | --- |
| Involvement of toxic metabolite derived from omega-6 PUFA | Exogenous omega-6 PUFAs sensitizes cells to ferroptosis (Doll et al., 2017). DGLA induces ferroptosis in HT-1080 cells (Figure 3). | Germ cell death induced by dietary omega-6 PUFAs DGLA and AA (Figs. 2, 3, 5 from (Watts and Browse, 2006)). |
| Involvement of oxidized lipids | Oxidized AA-PE and Ada-PE cause cell death (Kagan et al., 2017). | CYP450 produces toxic epoxides derived from DGLA (Figs. 2, 3, 4 from (Deline et al., 2015)). |
| Glutathione and oxidative stress resistance pathways (NRF2) protect from ferroptosis | Involvement of glutathione and GPX4 (Ingold et al., 2018; Seiler et al., 2008; Yang et al., 2014). Nrf2 protects cells from ferroptosis (Fan et al., 2017; Sun et al., 2016). | Mutations in glutathione synthesis genes sensitizes worms to DGLA-induced sterility. SKN-1 (NRF2) protects against cell death (Figs. 1, 2, 3, and 4 from (Webster et al., 2013)). RNAi knockdown of *gpx* genes sensitizes worms to DGLA-induced sterility (Figure 1F). |
| Apoptotic pathway not required | Caspase independence of ferroptosis (Dixon et al., 2012; Dolma et al., 2003) | DGLA-induced cell death occurs in apoptosis-defective mutants (Fig. 5 from (Webster et al., 2013), Figure 1A). |
| Specific inhibition by ferrostatin-1, iron chelators, and antioxidants. | Inhibition by ferrostatin-1 and iron chelators (Dixon et al., 2012; Skouta et al., 2014). Inhibition by antioxidants (Dixon et al., 2012; Matsushita et al., 2015; Yang and Stockwell, 2008, 2016) | DGLA-induced germ cell death is suppressed by ferrostatin-1, vitamin E, and iron chelators (Figure 1A, B, C, D, E, H). |
| Promotion of cell death by NADPH oxidase | NOX inhibitors suppress ferroptosis (Dixon et al., 2012) | *bli-3* mutants are resistant to DGLA-induced sterility (Figure 1G). |
| MUFAs protect from ferroptosis | MUFAs protect HT-1080 cells from erastin-induced ferroptosis (Magtanong et al., 2019) | Exogenous and endogenous MUFAs protect germ cells from DGLA-induced cell death (Figure 2). |

In *C. elegans* germ cell death can be induced by dietary DGLA, and to a lesser extent dietary AA, but not by linoleic acid (LA), or omega-3 PUFAs such as linolenic acid (ALA) or EPA (Watts and Browse, 2006). In mammalian cells, oxidized omega-6 PUFAs, but not omega-3 PUFAs, are specifically associated with the induction of ferroptosis (Doll et al., 2017; Kagan et al., 2017). Likewise, in mammalian HT-1080 cells, we find that DGLA can induce ferroptosis while omega-3 PUFAs like alpha-linolenic acid do not (Magtanong et al., 2019). These findings support the notion that ferroptosis is promoted by enzymatically-catalyzed selective initiation of lipid peroxidation (Deline et al., 2015; Kagan et al., 2017; Yang et al., 2016), rather than non-specific oxidation of any available PUFA. The role of ether lipids has not previously been explored in the context of ferroptosis, and our results suggest that these species may normally protect cells from ferroptosis, with the vinyl ether bonds possibly acting as oxidative sinks that protect nearby PUFA acyl chains from oxidation (Engelmann, 2004; Shi et al., 2016; Wallner and Schmitz, 2011). Humans born with defective ether lipid synthesis enzymes have severe growth and mental deficiencies and most do not survive beyond childhood (Wanders and Waterham, 2006). Our *C. elegans* findings raise the possibility that the failure to combat cellular oxidative stress, and perhaps the failure to prevent excessive ferroptotic cell death after exposure to dietary PUFAs, may contribute to the observed developmental defects.

Mechanistically, DGLA uptake is associated with a broad spectrum of changes in the lipidome of mammalian cells, with pro-ferroptotic doses of DGLA correlating with increased incorporation into phosphatidylcholines rather than phosphatidylethanolamines. We infer that exogenous DGLA may be initially buffered via incorporation into neutral lipids, but that at high levels this buffering capacity is exceeded, resulting in incorporation into membrane phospholipids. DGLA also displaces non-oxidizable MUFAs from these and other phospholipids, which may also enhance the overall oxidizability of the membrane (Magtanong et al., 2019). Notably, in both *C. elegans* and mammalian cells, DGLA uptake induced ferroptosis under conditions in which the glutathione peroxidase and other cellular antioxidant systems are not inactivated, but rather presumably overcome by an excess of oxidized lipids that exceed the capacity of these systems. This likely accounts for why high levels of DGLA incorporation are required to induce ferroptosis as a single agent. One possibility suggested by this work is that combining exogenous DGLA with small molecule inducers of ferroptosis may represent an especially effective means of inducing ferroptosis in cancer cells, as well as in other sensitive cell types in vivo.

## ACKNOWLEDGEMENTS

Some nematode strains were provided by the Caenorhabditis Genetics Center, funded by NIH Office of Research Infrastructure Programs (P40 OD010440). The GA912 *ftn-1(ok3625)* strain was a gift from Dr. David Gems (University College London, London, UK). The OB266 *bli-3(im10)* strain was a gift from Dr. Danielle Garsin (University of Texas Health Science Center, Houston, Texas, USA). Funding to JLW provided by the Farrell Memorial Fund, the Stanly Adler Research fund, and the Donald R. Weldin research fund. Funding to S.J.D. provided by the NIH (1R01GM122923).

## AUTHOR CONTRIBUTIONS

Conceptualization: J.L.W. and S.J.D. Methodology and Investigation: M.A.P, L.M.; Writing, editing, and reviewing: M.A.P., L.M., S.J.D., J.L.W.; Funding acquisition and Supervision: J.L.W. and S.J.D.

## DECALRATION OF INTERESTS

S.J.D. is a member of the scientific advisory board of Ferro Therapeutics.

## Supplemental Experimental Procedures

### Contact for Reagent and Resource Sharing

Further information and requests for resources and reagents should be directed to and will be fulfilled by the Lead Contact, Jennifer Watts.

## EXPERIMENTAL MODEL AND SUBJECT DETAILS

### *C. elegans* strains and maintenance

Nematode stocks were maintained on nematode growth media (NGM) plates seeded with bacteria (*E. coli* OP50) at 20°C. The following strains/alleles used in this study were obtained from the *Caenorhabditis* Genetics Center (CGC): N2 Bristol (wild-type), CB767 *bli-3(e767)*, MT1522 *ced-3(n717)*. The GA912 *ftn-1(ok3625)* strain was a gift from Dr. David Gems (University College London, London, UK). The OB266 *bli-3(im10)* strain was a gift from Dr. Danielle Garsin (University of Texas Health Science Center, Houston, Texas, USA).

### Cell lines and chemicals

HT-1080 cells (Cat# ATCC CCL-121) were purchased from ATCC, expanded for one passage, aliquoted and frozen down at −140°C until use. HT-1080^N^ cells (gender: male) were described previously (Forcina et al., 2017). Erastin2 (compound 35MEW28 in (Dixon et al., 2014)) was synthesized by Acme Bioscience (Palo Alto, CA). Dimethyl sulfoxide (DMSO, Cat# 276855) and ferrostatin-1 (Cat# SML0583) were from Sigma-Aldrich (St. Louis, MO). Ethanol (Cat# AX0441-3) was from EMD Millipore (Billerica, MA). Dihomo-γ-linolenic acid (DGLA, Cat# 90230) was from Cayman Chemical (Ann Arbor, MI). SYTOX Green (Cat# S7020) was from Molecular Probes (Eugene, OR). All compounds were stored at −20°C.

## METHODS DETAILS

### Fatty acid supplementation for *C. elegans* sterility assay

The fatty acid supplemented media was previously described (Deline et al., 2013). To NGM media, 0.1% Tergitol NP40 (Sigma Chemicals Cat#NP40S) and the sodium salt of dihomo-gamma-linolenic acid (DGLA, 20:3n-6) (NuChek Prep, Inc. Cat#S-1143) or oleic acid (OA, 18:1n-9) (NuChek Prep, Inc. Cat#S-1120) were added at various concentrations ranging from 0.1mM to 0.3mM DGLA. *E. coli* OP50 food source were seeded plates 3 days prior to plating *C. elegans* L1 synchronized larvae. Adult worms were scored for sterility at 72-96 hours where sterility was determined by scoring worms with absences of embryos as indicated by an empty uterus with light microscopy. The sterility tests were scored and averaged over five individual plates of each genotype, each consisting of 50 nematodes. Sterility was compared between control populations and experimental populations using a two-tailed student’s t-test. Exogenous fatty acid uptake of nematodes was confirmed through direct transesterification of nematode populations using 2.5% H_2_SO_4_ for one hour at 70°C to generate fatty acid methyl ester (FAMEs). FAMES were extracted with hexane for gas chromatography/mass spectrometry (GC/MS) analysis (Watts and Browse, 2002).

### Ferrostatin-1 and DGLA supplementation

DGLA plates were made ahead of time by adding various concentrations to molten NGM agar prior to plating. OP50 was plated and allowed to dry for two days prior to plating drugs and worms. The ferrostatin-1 (Fer-1) (Sigma-Aldrich Cat#SML0583) was dissolved in dimethyl sulfoxide (DMSO, Cat# 276855) and diluted to the appropriate concentrations in 10% DMSO prior to plating onto the OP50 food source shortly. The Fer-1 solution or control DMSO was allowed to absorb into the plates for 20-30 minutes before adding L1 stage nematodes.

### Trolox and 2,2’-bipyridine (BP) DGLA supplementation

2,2-bipyridiyl (BP) (Sigma-Aldrich Cat# D216305) (75μm) or Trolox (Cayman Chemicals Cat#10011659) were added directly to NGM agar. Either the drug or the equivalent volume amount of vehicle, DMSO, were added to molten along with various concentrations of DGLA into NGM agar prior to pouring into plates.

### RNA mediated interference (RNAi) of glutathione peroxidase genes on DGLA

RNAi by feeding on DGLA supplemented plates was performed as previously described (Deline et al., 2013). *E. coli* HT115 expressing the L4440 empty vector control or the *gpx* RNAi clones were obtained from the Ahringer Library (Source Biosource) and verified by sequencing (Kamath et al., 2003).

### DAPI staining for germ cell visualization

To visualize germ cells after feeding (either DGLA / DMSO, DGLA / 250μM Fer-1, or DGLA/1mM Trolox) worms were washed off with 1mL of M9 buffer and dispensed onto a watch glass. The majority of the M9 buffer was blotted dry, and worms were then fixed and stained with ethanol containing 0.2μg/mL 2’,6-diamidino-2-phenylindole (DAPI) (Fisher Scientific, Cat#D1306) and allowed to fix and stain for 10 minutes. Worms were placed on a coverslip with a droplet of water and adhered to a 2% agar pad containing slide. Images were acquired using an Olympus BX53 microscope (Olympus, Shinjuku, Tokyo, Japan) with a 10X objective lens.

### Mammalian cell death experiments

The day before the experiment, 5,000 HT-1080^N^ cells/well were seeded into a 96-well plate (Cat# 3904, Corning). The next day, the cell culture medium was replaced with DMSO (vehicle) or Fer-1 (1 μM) and ethanol (vehicle) or DGLA (2-fold, 10 point dose response series, starting at 500 μM). SYTOX Green (20 nM) was included in all wells. Erastin2 (1 μM) treatment was a positive control for ferrostatin-1-dependent inhibition of ferroptosis. Cells were imaged on the Essen IncuCyte Zoom (Ann Arbor, MI) and analyzed as described (Forcina et al., 2017). Three independent biological experiments were performed for each condition.

### C11 581/591 BODIPY imaging

The day before the experiment, 125,000 HT-1080 cells/well were seeded into 6-well plates (Cat# 3516, Corning) with one 22 mm^2^ no. 1.5 glass coverslip in each well. The next day, the cells were treated with vehicle (ethanol) or DGLA (500 μM) and DMSO or ferrostatin-1 (1 μM) in HT-1080 growth medium at 37°C for 8 h. After 8 h, the medium was removed and the cells were treated with C11 BODIPY 581/591 (Cat# D3861, Molecular Probes, Eugene, OR; final concentration = 5 μM) and Hoechst (Cat# H1399, Molecular Probes; final concentration = 1 μg/mL) dissolved in HBSS and incubated at 37°C for 10 min. After 10 min, the C11 BODIPY 581/591/Hoechst mixture was removed and fresh HBSS was added to the cells. Each cover slip was removed and mounted in 25 μL HBSS onto a glass microscope slide. Cells were imaged using a Zeiss Axio Observer microscope with a confocal spinning-disk head (Yokogawa, Tokyo, Japan), PlanApoChromat 63×/1.4 NA oil immersion objective, and a Cascade II:512 electron-multiplying (EM) CCD camera (Photometrics, Tucson, AZ). Images were processed in ImageJ 1.48v. Imaging was performed on two independent biological replicates per treatment condition.

### Mammalian cell lipid analysis

The day before the experiment, 2 × 10^6^ HT-1080 cells/plate were seeded into 10 cm dishes (Cat# CC7682-3394, USA Scientific, Ocala, FL). Four 10 cm dishes were seeded for each treatment. The next day, cells were treated with ethanol (vehicle) or dihomo-gamma-linoleic acid (Cat# 90230, Cayman Chemical, 250 μM or 500 μM). After 6 h, the media was removed and each plate was washed once with 3 ml 1× phosphate buffered saline, pH 7.4 (PBS, Cat# 10010023, Gibco). After washing, 2 mL of 1× PBS was added to each plate. Cells were scraped off each plate and transferred via glass Pasteur pipette to a conical bottomed glass tube (Cat# 05-569-2, Fisher Scientific), resulting in ∼8 mL total cell suspension per treatment. 0.5 mL was removed for protein determination. Each cell suspension (for both lipidomics and protein determination) was centrifuged (1000 rpm, 5 min, room temperature), the supernatant was removed using a glass Pasteur pipette, and the cell pellet was immediately stored at −80 °C, prior to further analysis. Protein concentration was determined using a BCA assay kit (Pierce). The remaining cell pellet was extracted over night at −20°C in 5 ml of ice-cold 5 ml chloroform:methanol (1:1). A solution of 0.2M H_3_PO_4_, 1M KCl was added to samples, which resulted in phase separation of the organic and aqueous phase. The organic phase was removed and dried under argon, then resuspended in chloroform. Samples were loaded in triplicate with authentic standards and TLC plates were developed in a solvent system consisting of chloroform:methanol:water:acetic acid (65:43:3:2.5). Lipids were visualized under UV light after spraying the plate with 0.005% primuline, and spots corresponding to phosphatidylcholine (PC), phosphatidylethanolamine (PE) and neutral lipids were scraped, spiked with a known standard (15:0), and transesterified in 2.5% H_2_SO_4_ to generate FAMES for GC/MS analysis to determine the fatty acid composition of each lipid class (Watts and Browse, 2002). Three biological replicates were used for TLC analysis.

