## Supplemental Table 1 for "Dietary Induction and Modulation of Ferroptosis in *Caenorhabditis elegans*"

Supplementary Table 1

Sterility Data and DGLA update data relating to Figures 1 and 2 N= number of plates of 50 worms each

Data for Figure 1A

| Treatment (WT) | % Sterility | % DGLA in worm lipids | N |
|---|---|---|---|
| 0mM DGLA / DMSO | 0% | 3.57% | 6 |
| 0mM DGLA / 125µM Fer-1 | 0% | 3.49% | 6 |
| 0mM DGLA / 250µM Fer-1 | 0% | 3.66% | 6 |
| 0.1mM DGLA / DMSO | 52% | 7.96% | 6 |
| 0.1mM DGLA / 125µM Fer-1 | 23% | 8.40% | 6 |
| 0.1mM DGLA / 250µM Fer-1 | 15% | 8.51% | 6 |
| 0.13mM DGLA / DMSO | 75% | 9.55% | 6 |
| 0.13mM DGLA / 125µM Fer-1 | 38% | 10.44% | 6 |
| 0.13mM DGLA / 250µM Fer-1 | 32% | 10.48% | 6 |

| Treatment (*ced-3* ) | % Sterility | % DGLA in worm lipids | N |
|---|---|---|---|
| 0mM DGLA / DMSO | 0% | 3.62% | 6 |
| 0mM DGLA / 125µM Fer-1 | 0% | 3.67% | 6 |
| 0mM DGLA / 250µM Fer-1 | 0% | 3.75% | 6 |
| 0.1mM DGLA / DMSO | 26% | 8.75% | 6 |
| 0.1mM DGLA / 125µM Fer-1 | 3% | 9.90% | 6 |
| 0.1mM DGLA / 250µM Fer-1 | 1% | 8.72% | 6 |
| 0.13mM DGLA / DMSO | 42% | 10.73% | 6 |
| 0.13mM DGLA / 125µM Fer-1 | 16% | 11.17% | 6 |
| 0.13mM DGLA / 250µM Fer-1 | 6% | 10.96% | 6 |

Data for Figure 1D

| Treatment (WT) | % Sterility | % DGLA in worm lipids | N |
|---|---|---|---|
| 0mM DGLA / DMSO (0.5) | 0% | 5.54% | 5 |
| 0mM DGLA / DMSO (1) | 0% | 4.50% | 5 |
| 0mM DGLA / 0.5mM Trolox | 0% | 4.41% | 5 |
| 0mM DGLA / 1mM Trolox | 0% | 4.22% | 5 |
| 0.1mM DGLA / DMSO (0.5) | 12% | 9.53% | 5 |
| 0.1mM DGLA / DMSO (1) | 13% | 8.60% | 5 |
| 0.1mM DGLA / 0.5mM Trolox | 0% | 8.30% | 5 |
| 0.1mM DGLA / 1mM Trolox | 0% | 8.82% | 5 |
| 0.15mM DGLA / DMSO (0.5) | 67% | 9.41% | 5 |
| 0.15mM DGLA / DMSO (1) | 71.2% | 10.09% | 5 |
| 0.15mM DGLA / 0.5mM Trolox | 37.2% | 9.44% | 5 |
| 0.15mM DGLA / 1mM Trolox | 25.2% | 9.49% | 5 |

Data for Figure 1G

| Treatment | % Sterility | % DGLA in worm lipids | N |
|---|---|---|---|
| 0mM DGLA N2 | 0.0% | 3.67% | 5 |
| 0mM DGLA bli-3(e767) | 0.0% | 3.27% | 5 |
| 0mM DGLA bli-3(im10) | 0.0% | 3.49% | 5 |
| 0.15mM DGLA N2 | 62.4% | 8.33% | 5 |
| 0.15mM DGLA bli-3(e767) | 7.2% | 11.47% | 5 |
| 0.15mM DGLA bli-3(im10) | 4.4% | 11.66% | 5 |
| 0.3mM DGLA N2 | 90.8% | 14.05% | 5 |
| 0.3mM DGLA bli-3(e767) | 15.2% | 17.40% | 5 |
| 0.3mM DGLA bli-3(im10) | 9.2% | 15.35% | 5 |

Data for Figure 1F

| Treatment | % Sterility | % DGLA in worm lipids | N |
|---|---|---|---|
| 0mM DGLA EV | 0.0% | 4.33% | 5 |
| 0mM DGLA gpx-1 | 0.0% | 3.78% | 5 |
| 0mM DGLA gpx-6 | 0.0% | 4.03% | 5 |
| 0mM DGLA gpx-7 | 0.0% | 3.73% | 5 |
| 0.2mM DGLA EV | 24.8% | 10.24% | 5 |
| 0.2mM DGLA gpx-1 | 92.0% | 9.59% | 5 |
| 0.2mM DGLA gpx-6 | 82.0% | 9.69% | 5 |
| 0.2mM DGLA gpx-7 | 98.0% | 9.52% | 5 |
| 0.4mM DGLA EV | 73.6% | 11.34% | 5 |
| 0.4mM DGLA gpx-1 | 98.8% | 11.91% | 5 |
| 0.4mM DGLA gpx-6 | 100.0% | 10.63% | 5 |
| 0.4mM DGLA gpx-7 | 98.4% | 10.12% | 5 |

Data for Figure 1H

| Treatment | % Sterility | % DGLA in worm lipids | N |
|---|---|---|---|
| 0mM DGLA N2 DMSO | 0.0% | 3.81% | 6 |
| 0mM DGLA ftn-1 DMSO | 0.0% | 3.80% | 6 |
| 0mM DGLA N2 75BP | 0.0% | 6.87% | 6 |
| 0mM DGLA ftn-1 75BP | 0.0% | 9.31% | 6 |
| 0.1mM DGLA N2 DMSO | 49.3% | 8.87% | 6 |
| 0.1mM DGLA ftn-1 DMSO | 70.0% | 9.09% | 6 |
| 0.1mM DGLA N2 75BP | 5.3% | 15.79% | 6 |
| 0.1mM DGLA ftn-1 75BP | 29.3% | 14.33% | 6 |
| 0.15mM DGLA N2 DMSO | 76.0% | 12.08% | 6 |
| 0.15mM DGLA ftn-1 DMSO | 88.7% | 10.77% | 6 |
| 0.15mM DGLA N2 75BP | 18.0% | 15.78% | 6 |
| 0.15mM DGLA ftn-1 75BP | 33.3% | 19.49% | 6 |

Data for Figure 2C

| Treatment | % Sterility | % DGLA in worm lipids | N |
|---|---|---|---|
| 0mM DGLA N2 | 0.0% | 3.81% | 5 |
| 0mM DGLA ads-1 | 0.0% | 4.40% | 5 |
| 0.1mM DGLA N2 | 44.4% | 9.55% | 5 |
| 0.1mM DGLA ads-1 | 92.0% | 7.58% | 5 |
| 0.15mM DGLA N2 | 68.8% | 11.94% | 5 |
| 0.15mM DGLA ads-1 | 98.0% | 8.74% | 5 |

Data for Figure 2D

| Treatment | % Sterility | % DGLA in worm lipids | N |
|---|---|---|---|
| 0mM DGLA/DMSO N2 | 0.0% | 3.87% | 5 |
| 0mM DGLA/DMSO ads-1 | 0.0% | 3.63% | 5 |
| 0.05mM DGLA/DMSO N2 | 0.4% | 6.79% | 5 |
| 0.05mM DGLA/125uM Fer-1 N2 | 0.0% | 6.80% | 5 |
| 0.05mM DGLA/250uM Fer-1 N2 | 0.0% | 6.81% | 5 |
| 0.05mM DGLA/DMSO ads-1 | 64.4% | 4.54% | 5 |
| 0.05mM DGLA/125uM Fer-1 ads-1 | 48.4% | 4.70% | 5 |
| 0.05mM DGLA/250uM Fer-1 ads-1 | 29.2% | 4.67% | 5 |
| 0.075mM DGLA/DMSO N2 | 6.8% | 7.83% | 5 |
| 0.075mM DGLA/125uM Fer-1 N2 | 1.6% | 7.32% | 5 |
| 0.075mM DGLA/250uM Fer-1 N2 | 0.8% | 7.55% | 5 |
| 0.075mM DGLA/DMSO ads-1 | 80.8% | 4.92% | 5 |
| 0.075mM DGLA/125uM Fer-1 ads-1 | 67.6% | 5.03% | 5 |
| 0.075mM DGLA/250uM Fer-1 ads-1 | 59.6% | 5.20% | 5 |
