## Supplemental Table 2 for "Dietary Induction and Modulation of Ferroptosis in *Caenorhabditis elegans*"

Supplementary Table 2
GC/MS data of fatty acid composition relating to Figure 2

Data for Figure 2A, 2B, and 2C

| Treatment | % Sterility | N | 14:0 | 15iso | 16:0 | 17iso | 16:1 | 17Δ | 18:0 | 18:1n-9 | 18:1n-7 | 18:2 | 19Δ | 20:3 | 20:4 | 20:4n-3 | 20:5 |
|---|---|---|---|---|---|---|---|---|---|---|---|---|---|---|---|---|---|
| 0mM DGLA/0mM OA N2 | 0.0% | 5 | 0.98% | 3.06% | 3.86% | 2.45% | 1.01% | 17.38% | 4.96% | 3.22% | 11.54% | 3.64% | 24.86% | 3.76% | 0.74% | 5.01% | 10.97% |
| 0mM DGLA/0.1mM OA N2 | 0.0% | 5 | 0.82% | 3.50% | 4.18% | 3.37% | 1.40% | 10.51% | 5.41% | 6.09% | 11.10% | 4.69% | 23.85% | 3.84% | 0.77% | 5.79% | 11.76% |
| 0mM DGLA/0.2mM OA N2 | 0.0% | 5 | 0.76% | 3.67% | 4.00% | 3.63% | 1.40% | 8.53% | 5.22% | 7.11% | 10.28% | 4.91% | 25.70% | 3.76% | 0.68% | 5.67% | 11.82% |
| 0mM DGLA/0.3mM OA N2 | 0.0% | 5 | 0.75% | 3.75% | 3.89% | 3.91% | 1.34% | 6.61% | 5.46% | 8.47% | 9.91% | 5.10% | 25.55% | 3.65% | 0.66% | 5.93% | 12.20% |
| 0.1mM DGLA/0mM OA N2 | 44.4% | 5 | 1.64% | 3.49% | 4.61% | 1.97% | 2.09% | 9.61% | 5.67% | 4.79% | 7.51% | 5.29% | 28.82% | 6.60% | 0.85% | 6.52% | 8.32% |
| 0.1mM DGLA/0.1mM OA N2 | 0.0% | 5 | 1.10% | 4.44% | 4.99% | 3.75% | 1.68% | 9.53% | 6.38% | 5.71% | 10.76% | 5.05% | 15.49% | 5.86% | 0.90% | 7.61% | 13.52% |
| 0.1mM DGLA/0.2mM OA N2 | 0.0% | 5 | 0.86% | 3.93% | 4.20% | 3.67% | 1.40% | 7.13% | 5.54% | 5.73% | 8.73% | 5.27% | 26.77% | 4.96% | 0.69% | 6.41% | 11.75% |
| 0.1mM DGLA/0.3mM OA N2 | 0.0% | 5 | 0.81% | 4.01% | 3.93% | 3.82% | 1.42% | 6.70% | 5.18% | 6.50% | 8.52% | 5.17% | 26.72% | 4.54% | 0.68% | 6.85% | 12.34% |
| 0.2mM DGLA/0mM OA N2 | 66.8% | 5 | 2.01% | 3.61% | 5.01% | 1.79% | 2.51% | 7.75% | 6.76% | 4.87% | 6.60% | 5.91% | 20.49% | 11.34% | 1.15% | 9.17% | 8.59% |
| 0.2mM DGLA/0.1mM OA N2 | 0.0% | 5 | 0.95% | 3.78% | 4.57% | 3.29% | 1.58% | 8.07% | 5.40% | 4.91% | 8.36% | 5.65% | 24.72% | 6.64% | 0.79% | 7.38% | 10.91% |
| 0.2mM DGLA/0.2mM OA N2 | 0.0% | 5 | 0.84% | 3.93% | 4.24% | 3.52% | 1.53% | 7.20% | 5.16% | 5.68% | 8.07% | 5.25% | 27.46% | 5.94% | 0.70% | 6.98% | 10.81% |
| 0.2mM DGLA/0.3mM OA N2 | 0.0% | 5 | 0.78% | 4.04% | 4.10% | 3.82% | 1.49% | 6.42% | 5.39% | 6.72% | 8.13% | 6.77% | 26.20% | 5.75% | 0.74% | 6.57% | 10.33% |
| 0.3mM DGLA/0mM OA N2 | 94.8% | 5 | 2.01% | 3.59% | 5.12% | 1.68% | 2.44% | 7.49% | 6.71% | 4.87% | 5.97% | 5.16% | 22.36% | 13.48% | 1.09% | 8.80% | 6.95% |
| 0.3mM DGLA/0.1mM OA N2 | 13.2% | 5 | 0.99% | 3.87% | 4.46% | 3.08% | 1.83% | 7.96% | 5.40% | 5.08% | 7.89% | 6.41% | 22.24% | 8.55% | 0.86% | 8.46% | 10.04% |
| 0.3mM DGLA/0.2mM OA N2 | 2.4% | 5 | 0.85% | 3.84% | 4.29% | 3.38% | 1.54% | 6.41% | 5.54% | 5.37% | 7.41% | 6.03% | 25.24% | 7.65% | 0.74% | 8.34% | 10.48% |
| 0.3mM DGLA/0.3mM OA N2 | 1.6% | 5 | 0.79% | 3.94% | 4.10% | 3.61% | 1.50% | 5.88% | 5.40% | 5.87% | 7.27% | 6.59% | 25.42% | 7.59% | 0.71% | 8.24% | 10.31% |
| 0mM DGLA/0mM OA fat-2 | 25.2% | 5 | 0.88% | 2.81% | 1.78% | 4.50% | 3.69% | 17.89% | 2.14% | 20.56% | 24.20% | - | 18.12% | - | - | - | 1.14% |
| 0mM DGLA/0.1mM OA fat-2 | 36.0% | 5 | 0.79% | 3.03% | 1.69% | 5.27% | 3.76% | 11.39% | 2.12% | 23.46% | 24.22% | - | 21.01% | - | - | - | 1.06% |
| 0mM DGLA /0.2mM OA fat-2 | 41.2% | 5 | 0.79% | 3.08% | 1.68% | 5.50% | 3.60% | 9.67% | 2.06% | 24.14% | 23.46% | - | 23.10% | - | - | - | 1.02% |
| 0mM DGLA/0.3OA fat-2 | 50.0% | 5 | 0.78% | 3.11% | 1.60% | 5.64% | 3.44% | 8.12% | 2.06% | 25.20% | 23.15% | - | 23.65% | - | - | - | 1.19% |
| 0.1mM DGLA/0mM OA fat-2 | 3.2% | 5 | 1.15% | 4.23% | 4.07% | 2.83% | 1.86% | 11.37% | 5.90% | 20.23% | 9.23% | - | 18.21% | 4.79% | 0.93% | 4.15% | 8.86% |
| 0.1mM DGLA/0.1mM OA fat-2 | 0.0% | 5 | 0.95% | 3.62% | 2.13% | 4.46% | 3.04% | 8.00% | 3.08% | 23.28% | 18.12% | - | 24.24% | 0.91% | 0.19% | 1.59% | 4.60% |
| 0.1mM DGLA/0.2mM OA fat-2 | 0.0% | 5 | 0.93% | 3.57% | 1.81% | 5.02% | 3.21% | 7.32% | 2.59% | 24.09% | 20.13% | - | 24.32% | 0.68% | 0.00% | 1.03% | 3.48% |
| 0.1mM DGLA/0.3mM OA fat-2 | 0.0% | 5 | 0.98% | 3.88% | 1.93% | 5.84% | 3.49% | 5.60% | 3.38% | 26.91% | 22.46% | - | 20.12% | 0.58% | 0.29% | 0.86% | 2.11% |
| 0.2mM DGLA/0mM OA fat-2 | 0.0% | 5 | 1.09% | 4.05% | 4.12% | 2.47% | 1.76% | 10.11% | 5.92% | 17.78% | 8.17% | - | 16.80% | 8.93% | 1.34% | 5.52% | 9.87% |
| 0.2mM DGLA/0.1mM OA fat-2 | 0.0% | 5 | 0.88% | 3.92% | 3.02% | 3.75% | 2.25% | 8.75% | 4.48% | 23.93% | 11.75% | - | 24.64% | 1.95% | 0.41% | 2.29% | 6.05% |
| 0.2mM DGLA/0.2mM OA fat-2 | 48.0% | 5 | 0.89% | 3.62% | 2.44% | 4.15% | 2.55% | 7.35% | 3.88% | 23.46% | 15.04% | - | 24.87% | 1.43% | 0.35% | 2.13% | 6.07% |
| 0.2mM DGLA/0.3mM OA fat-2 | 68.8% | 5 | 0.88% | 3.77% | 1.96% | 4.75% | 2.86% | 6.71% | 2.93% | 24.78% | 17.26% | - | 24.68% | 1.04% | 0.25% | 1.73% | 4.79% |
| 0.3mM DGLA/0mM OA fat-2 | 0.8% | 5 | 1.01% | 3.90% | 4.14% | 2.33% | 1.73% | 9.15% | 6.03% | 15.63% | 7.39% | - | 14.92% | 13.59% | 1.61% | 6.85% | 9.73% |
| 0.3mM DGLA/0.1mM OA fat-2 | 0.0% | 5 | 0.97% | 3.88% | 3.60% | 3.30% | 1.96% | 7.99% | 5.20% | 21.93% | 9.11% | - | 21.73% | 4.49% | 0.82% | 4.29% | 9.04% |
| 0.3mM DGLA/0.2mM OA fat-2 | 0.0% | 5 | 0.83% | 3.68% | 3.34% | 3.66% | 1.84% | 6.97% | 5.17% | 24.37% | 9.67% | - | 24.64% | 3.21% | 0.64% | 3.24% | 7.31% |
| 0.3mM DGLA/0.3mM OA fat-2 | 0.0% | 5 | 0.79% | 4.12% | 3.10% | 3.77% | 1.77% | 6.72% | 4.97% | 24.98% | 9.62% | - | 24.93% | 2.97% | 0.50% | 2.97% | 7.22% |

**abbreviations**

fatty acid nomenclature x:y(n-z) x= # of carbons, y= # of double bonds, n-z = location of terminal double bonds, z carbons from the methyl end

15:*iso* 13-methyltetradecanoic acid

17:*iso* 15-methylhexanoic acid

17Δ cis-9,10-methylenehexadecanoic acid

19Δ cis-11,12-methylene octadecanoic acid

- not detected (<0.25%)
